# A coordinated morphogenetic program drives rapid body plan transformation in a sponge

**DOI:** 10.64898/2026.04.23.719999

**Authors:** Océane Blard, Zac Pujic, Stefan Thor, Sandie M. Degnan, Bernard M. Degnan

## Abstract

Across the animal kingdom, metamorphosis transforms a free-swimming larva into a morphologically distinct juvenile, yet how conserved cellular processes are spatiotemporally coordinated to execute this rapid body plan switch remains poorly understood. Using the marine sponge *Amphimedon queenslandica* — a member of one of the earliest-diverging animal phyletic lineages — we characterise the cellular and morphogenetic events of the first six hours of metamorphosis. We show that metamorphosis proceeds through a tightly ordered sequence of events orchestrated by a stereotypic wave of epithelial infoldings that propagates from the basal to the apical pole within the first hour post-settlement. This wave acts as a morphogenetic pacemaker, spatially and temporally inducing subsequent cellular transitions, which include coordinated mucus secretion, cilia resorption, epithelial-mesenchymal transition (EMT), mesenchymal-epithelial transition (MET), transdifferentiation and targeted programmed cell death. Lineage tracing further reveals the stepwise transformation of larval cells at metamorphosis, with labelled epithelial flask cells transdifferentiating into internal archaeocyte stem cells via a transitory amoeboid cell state associated with EMT. These findings demonstrate that rapid metamorphosis comprises a spatially pre-patterned, stepwise program, and provide a cellular framework for understanding body plan transformations with broad implications for the evolution of metazoan metamorphosis.

## Introduction

Metamorphosis is the transformation of a larva into a morphologically and ecologically distinct juvenile, and is one of the most dramatic developmental events in the animal kingdom [1]. In marine invertebrates with pelagobenthic life cycles, this transition must be executed rapidly and reliably in response to environmental settlement cues, often producing a functional juvenile within hours to days [2]. Understanding how a metamorphosing postlarva achieves such a rapid and coherent body plan switch is a fundamental question in developmental and evolutionary biology.

The broad cellular vocabulary of metamorphosis is now well established. Across diverse animal phyla, the larva-to-juvenile transition involves a recurring set of cellular processes. These include: (i) directed epithelial reorganisation; (ii) epithelial-mesenchymal transition (EMT) and its reverse (mesenchymal-epithelial transition; MET); (iii) transdifferentiation of larval cell types into juvenile counterparts; (iv) differentiation of pluripotent stem cells alongside proliferation of potency-restricted progenitor populations; (v) programmed cell death (PCD); and (vi) coordinated secretory activity [2–9]. These processes are not bilaterian innovations, but rather are already present in cnidarians [10–12] and sponges [4, 13–29], indicating that the cellular toolkit of metamorphosis is ancestral to all animals. However, what remains poorly understood is how these cellular processes are spatiotemporally orchestrated to accomplish a specific body plan remodelling at the speed that metamorphosis often demands. This requires the simultaneous dismantling of structures essential for larval life, and the repurposing or making available existing cell populations to establish new juvenile tissue architectures and functions in a coordinated manner that maintains organismal integrity.

In bilaterian animals with complex larvae, the sheer number of cell types and tissues makes it challenging to resolve the sequence and interdependence of metamorphic events. Marine sponges offer an opportunity to meet this challenge. Their larvae are anatomically simple, typically comprising a small number of cell types arranged along a radially symmetric axis [21, 22, 24, 30, 31]. This comparative simplicity provides an opportunity for resolving the cellular logic of metamorphosis at a resolution that is difficult to achieve in more complex animals. The demosponge *Amphimedon queenslandica* is particularly well suited to such analysis because its larval cell types are defined molecularly and morphologically, and its metamorphosis is experimentally-inducible by contact with the coralline alga *Amphiroa fragilissima* and proceeds through a series of accurately stageable morphological forms [32–37]. This prior knowledge and predictability allow for cellular events to be assigned to defined temporal windows, their spatial distribution to be mapped, and their sequence to be causally ordered.

Here, we present an account of the cellular and morphogenetic events during the first six hours of *A. queenslandica* metamorphosis, with the goal of understanding not just which cellular processes are deployed but how their spatial organisation and temporal sequence initiate and drive a coherent postlarval morphogenetic program. We show that the program is organised around a stereotypic wave of epithelial infoldings that functions as a spatial and temporal pacemaker, inducing cellular transitions that follow in its wake. We use direct lineage tracing to demonstrate that larval sensory-secretory epithelial flask cells undergo a sequence of state transitions that lead to their transdifferentiation into archaeocyte stem cells, providing an example of how a specific larval cell type is repurposed during sponge metamorphosis. Together, our findings reveal how a small set of metazoan-conserved cellular mechanisms can be integrated into a rapid, coherent body plan switch, and establish a cellular framework for understanding the execution of morphogenetic programs in metamorphosis more broadly.

## Results

### A directional wave of epithelial infoldings establishes the spatial framework for metamorphosis

Within minutes of an *Amphimedon queenslandica* larva settling onto the coralline alga *Amphiroa fragilissima*, a rapid and spatiotemporally coordinated sequence of large-scale morphological changes is initiated (Fig. 1A-F and Supplementary videos 1-4). A key feature of this sequence is a wave of epithelial infoldings that propagates from the basal side of the postlarva – the region in contact with the alga and equivalent to the former larval anterior – toward the apical pole, reaching completion within approximately one hour post-settlement (hps) (Fig. 1A-D, G). This wave does not appear to be a passive mechanical consequence of substrate attachment, but rather it occurs in a stereotypical manner in all individuals, regardless of the geometry of the algal substrate or the precise region of larval contact (Fig. 1H-J). This is consistent with the wave being driven by an internally encoded program activated by the settlement cue.

**Figure 1.**
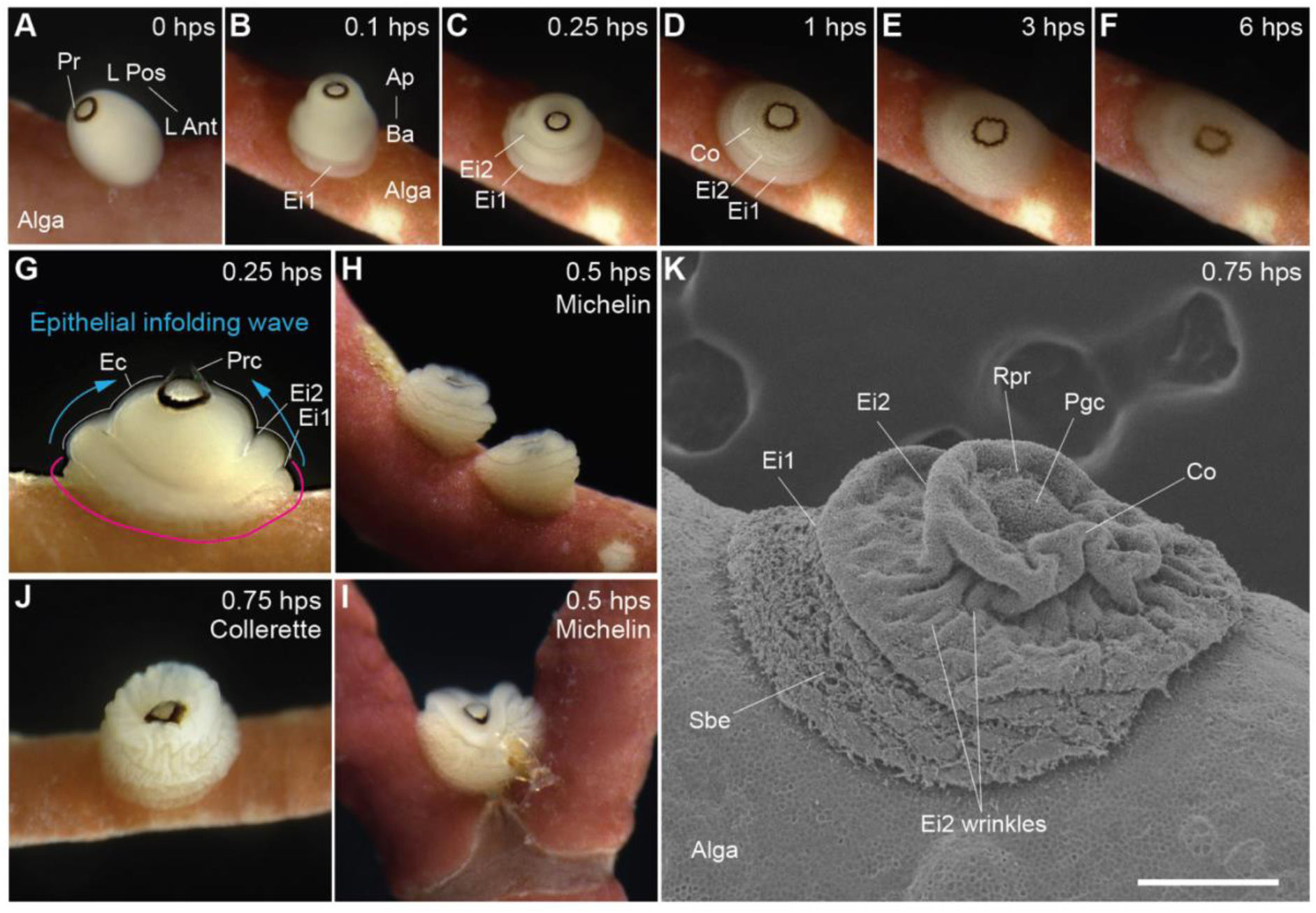
External characteristics of early metamorphosis of *A. queenslandica*. **(A-F)** Apicolateral views of postlarvae within the first six hours of settling on the coralline alga *A. fragilissima* (Alga). See Supplementary videos 1-4 for timelapses of the first hour of metamorphosis. **(A)** Competent larva contacting *A. fragilissima* by its anterior pole. **(B)** Newly settled postlarva within 5-10 min of initiating metamorphosis. The first epithelial infolding (Ei1) has commenced on the basal side (Ba) of the postlarva, and an intact pigment ring (Pr) is present on the apical side (Ap). **(C)** Postlarva at 0.25 hps undergoing progressive longitudinal epithelial infoldings with first and second infoldings (Ei2) present. **(D)** By 1 hps, the two infoldings have wrinkled and begun to collapse on each other with cells within Ei1 beginning to spread over the alga. **(E)** At 3 hps, the basal side of the postlarva continues to spread over the alga. **(F)** Flattening and spreading continue in the 6 hps postlarva as the pigment ring disintegrates and is covered over by egressing and migrating cells. **(G-J)** Examples of postlarvae in the first hour of metamorphosis. **(G)** Lateral view of a postlarva 15 min after settling with first and second epithelial infoldings present. The apicobasal morphogenetic wave is shown with blue arrows. Epithelial cilia are absent below the first infolding (pink line), and present above (Ec, white line) along with long pigment ring cilia (Prc). **(H, I)** 0.5 hps postlarvae with a Michelin-like shape from several epithelial infoldings. The infolding patterns of these three individuals illustrate the stereotypic nature of this process regardless of algal terrain or shape. The pigment ring begins to depress into the body. **(J)** Postlarva at 0.75 hps with a wrinkled body and an epithelial collerette that is around and extends above the pigment ring. As epithelial infoldings progress during the first 0.5 hps, the pigment ring subsides into the body, with the epithelium overlying the pigment ring wrinkling and forming a transient collerette at 0.75 hps. This structure collapses outward and repositions the pigment ring at the apex in 1 hps postlarvae. **(K)** Scanning electron micrograph of a 0.75 hps postlarva on *A. fragilissima*. First and second epithelial infoldings and an apical collarette are present. The disruption of larval epithelia integrity occurs below the first infolding in concert with the spreading of basal epithelial cells over the algae (Sbe). The pigment ring is still receding into the body (Rpr) and surrounded by a collerette-like epithelium. Posterior epithelial globular cells (Pgc) within the larval pigment ring are still present at this stage. Scale bar, 200 µm.

This wave of epithelial infolding manifests as follows. Within 15 minutes of settlement, the basal postlarval epithelium (formerly the larval anterior epithelium) compresses against the substrate, while the apical epithelium retains its smooth, ciliated larval morphology (Fig. 1B). A first deep longitudinal (annular) epithelial infolding then forms basally, just above the site of attachment (Fig. 1G). This first infolding is followed rapidly by a wave of successive longitudinal infoldings that form in basal-to-apical direction (Fig. 1C). This is followed by smaller latitudinal infoldings (epithelial wrinkles) that appear first at the base and advance toward the apex. By 0.5 hours post-settlement (hps), the postlarva has acquired a characteristic circumferential folded morphology (Michelin-like shape) with 3–4 longitudinal infoldings, and the larval posterior photosensory pigment ring has begun to subside into the postlarval body interior (Figs. 1H, I). Within further 15 min, these infoldings have wrinkled with the most apical forming a collerette-like shape (Fig. 1J, K). At this stage, the epithelial layer basal of the first infolding becomes disorganised and begins to spread over the algal surface. By 1 hps, the postlarval body has collapsed into a wrinkled, flattened ball, with the basal epithelium spreading further across the alga (Fig. 1D, K). Body flattening continues over the subsequent hours, with the spreading of postlarval cells over the algal substrate becoming more evident by 3 hps and substantially advanced by 6 hps (Fig. 1E, F).

The infolding wave appears not only as a morphogenetic event, but as a spatial pacemaker for the cellular transitions that follow. As each zone of the epithelium is overtaken by the advancing wave, it undergoes a series of local changes that are not yet occurring in zones apical to the wavefront, including ciliary resorption, globular cell secretion and PCD (see below for details). The basal-to-apical wave therefore imposes a temporal order on cellular events that is spatially restricted, acting to transform what might otherwise be a simultaneous, potentially disruptive wholesale reorganisation into a sequential, spatially coherent program.

### Changes in larval cell populations occur in a defined spatial and temporal sequence

Within the spatial framework established by the first infolding wave, different cell populations undergo state changes in a defined sequence. This does not appear to be solely a property of the wave itself, because different cell types respond differently to it, and some cell transitions occur independently of the wave’s immediate position. To understand this process, we first defined the roles and locations of larval cell types and tissues that rapidly transition in concert with, and independent of, this morphogenetic wave (Fig. 2).

**Figure 2.**
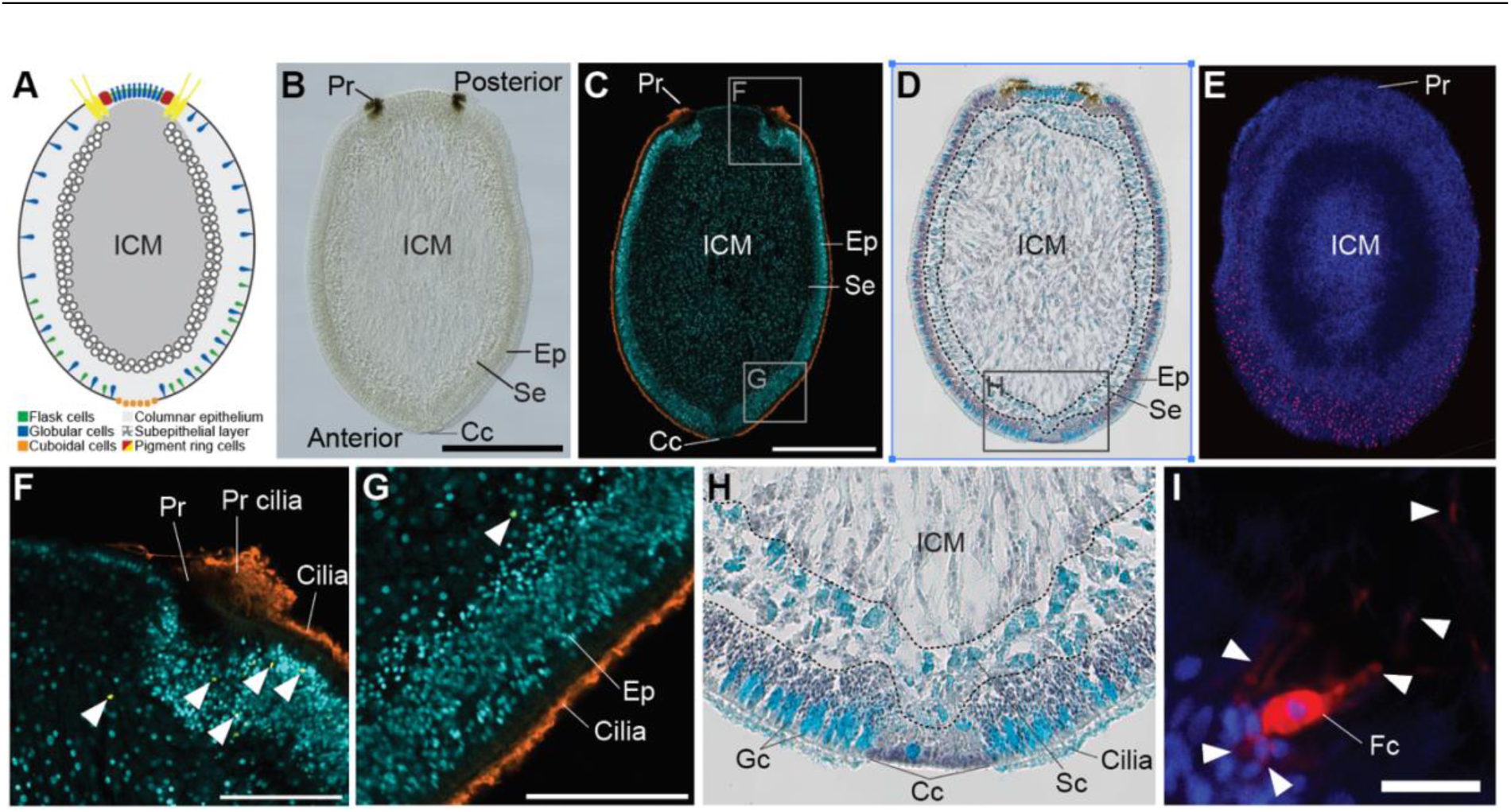
*A. queenslandica* larval cellular organisation and function. **(A)** Summary of major larval cell types and regions (adapted from [35]). **(B-E)** Larvae with anterior at the bottom, which will become the basal region of the metamorphosing postlarva; (B-D) longitudinal sections through the middle of the larva and (E) whole mount. **(B)** Differential interference contrast (DIC) micrograph showing pigmented cells of the posterior pigment ring (Pr), anterior cuboidal cells (Cc), epithelial (Ep) and subepithelial (Se) layers, and the inner cell mass (ICM). **(C)** Confocal micrograph with cilia labelled with anti-α-tubulin antibody (orange), fragmented nuclei of dead cells with TUNEL assay (yellow) and all nuclei with Hoechst (cyan). **(D)** DIC micrograph of larvae stained with alcian blue (cyan), and hematoxylin (purple, including nuclei). Cells and regions with high sulphated mucin are stained with alcian blue. Dashed lines demarcate boundaries between epithelial and subepithelial layers, and the subepithelial layer and the ICM. **(E)** Confocal micrographs showing flask cells in the epithelial layer labelled with CM-DiI (magenta) and all nuclei with Hoechst (cyan). **(F, G)** Magnifications of regions boxed in C showing numerous small nuclei of columnar epithelial cells that populate the surface of the larva (cyan) and surface cilia (orange). Pigment ring cilia are significantly longer than columnar epithelial cilia. Cells undergoing PCD are labelled in yellow and marked with white arrowheads. PCD is limited in the larva and largely localised to the vicinity of the pigment ring (F), although a few cells in the columnar epithelial layer are also apoptosing (G). **(H)** Magnification of the anterior region boxed in D showing that vesicles in epithelial globular cell (Gc) and subepithelial spherulous cells (Sc) contain sulphated mucins and that epithelial cilia are coated in mucins, likely from globular cell secretions (Supp Fig. 1). **(I)** Magnification of an epithelial flask cell (Fc) stained with DiI. White arrowheads point to apical and basal processes. Scale bars: B, C, 200 μm; E, 100 µm; F, G, 50 μm; I, 20 µm.

The larval epithelial layer that undergoes the basal-to-apical infoldings comprises of several morphologically discernible cell types: (i) thousands of tiny ciliated columnar epithelial cells dominate and cover the entire larval surface except at the most anterior and posterior regions, which are populated by cuboidal cells and a pigment ring, respectively (Fig. 2A-C, F, G); (ii) highly vacuolated globular cells rich in sulphated mucins are interspersed amongst the columnar epithelial cells as well as concentrated inside the pigment ring, and their putative progenitors – the spherulous cells in the subepithelial layer – are also enriched in these mucins (Fig. 2D, H and Supplementary Fig. 1) [38, 39]; (iii) flask cells with cellular processes that extend both externally and internally are interspersed in the epithelium but enriched anteriorly (Fig. 2E, I and Supplementary Fig. 1) [33]; and (iv) pigment and long-ciliated cells form the circular posterior pigment ring (Fig. 2B, C, F).

Together, these cell types function in the directional swimming of the larva, and in the detection and response to inductive settlement cues [14, 33, 40, 41]. The sulphated mucins appear to be released externally from the globular cells (Fig. 2H and Supplementary Fig. 1) and may serve to coat and potentially lubricate the cilia of the swimming larva [42, 43]. Globular cells, along with flask cells, which are often in proximity in the larval anterior pole (Supplementary Fig. 1), have a central role in detecting exogenous cues and promulgating inductive signals internally [33, 41], and pigment ring cells regulate larval phototaxis [40].

Programmed cell death (PCD), assessed by TUNEL labelling, is spatially restricted throughout early metamorphosis to the photosensory pigment ring and portions of the larval epithelium. Interestingly, several TUNEL-positive cells are present in the vicinity of the larval pigment ring (Fig. 2F), consistent with these cells precociously undergoing PCD, which is normally triggered early in metamorphosis (see Fig. 1A-F and below). Rarer PCD events also occur in the larval columnar epithelium (Fig. 2G).

### Ciliary resorption and re-elongation: transdifferentiation induced by the wave

The ciliated columnar cells of the larval epithelium, the beating cilia of which drive larval swimming behaviour, undergo a two-phase ciliary dynamic during early metamorphosis that is spatially coordinated by the first epithelial infolding wave. In zones of epithelium contacting the alga and already reached by the wave, cilia have ceased beating and are partially resorbed, while cilia in zones not yet overtaken by this wave remain intact and motile (Figs 1G, 3A-C). This wave of ciliary resorption proceeds in the same basal-to-apical direction as the infolding wave, coupling the mechanical reorganisation of the epithelium to the initiation of ciliary remodelling in a coordinated spatial sweep. Also coordinated with this wave appears to be a minor pruning of epithelial cells, with a subset undergoing PCD (Fig. 3B, C).

**Figure 3.**
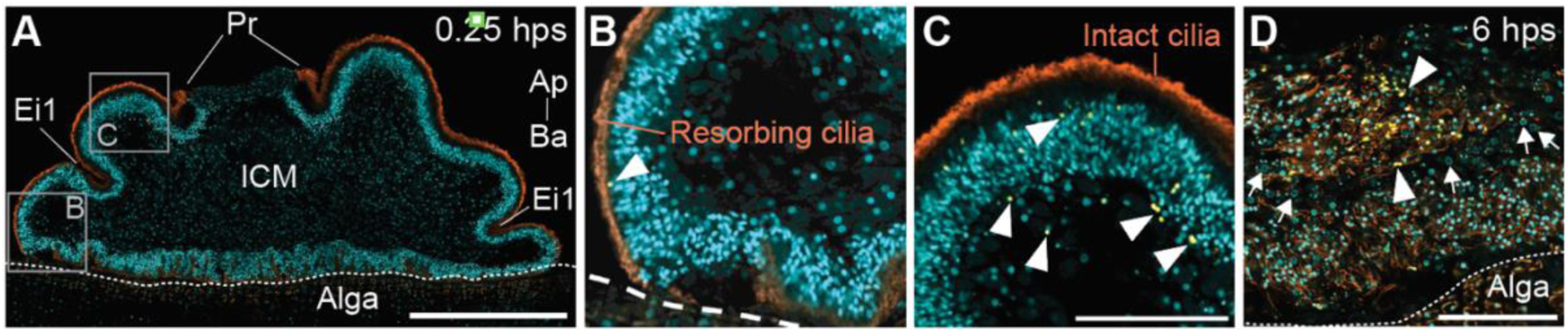
Resorption and re-elongation of epithelial cilia in the first hours of metamorphosis. Confocal micrographs of longitudinal sections of postlarvae settled on *A. fragilissima* (Alga); apical (Ap) – basal (Ba) axis shown. Cilia are labelled with an anti-α-tubulin antibody (orange), fragmented nuclei of dead cells labelled using TUNEL assay (yellow) and all nuclei labelled with Hoechst (cyan). **(A-C)** 0.25 hps postlarvae with epithelial cilia getting progressively resorbed. The region below the first epithelial infolding (Ei1) (B), cilia are closer to the epithelial layer (i.e. partially resorbed), compared to the cilia located above the infolding (C). **(D)** Internal view of 6 hps postlarva showing clusters of former larval epithelial cells that have undergone EMT and are re-elongating their cilia. These cells are now proximal to internal archaeocytes (white arrows pointing to their diagnostic larger nuclei with nucleoli). PCD continues to occur in larval epithelial cells (white arrowheads). Scale bars: A, 200 µm; C, D, 50 µm.

However, by 6 hps the cilia of the former larval epithelial cells, which have by now undergone EMT, have begun to re-elongate (Fig. 3D). This resorption-then-re-elongation sequence appears as a signature of the start of transdifferentiation of larval columnar epithelial cells into juvenile choanocytes, where locomotory cilia are first partially resorbed, and then elongated to form the choanocyte flagella [18]. That this process is already underway by 6 hps indicates that transdifferentiation is initiated early, within the first hours of metamorphosis, well before the formation of choanocytes and the juvenile body plan. The wave of epithelial infoldings therefore not only regulates the mechanics of ciliary resorption, but also triggers the onset of a transdifferentiation program that will ultimately produce the filter-feeding apparatus of the juvenile.

### Coordinated secretion in concert with the advancing wave

As the longitudinal infolding wave advances apically, globular cells progressively become unrecognisable in the zones of epithelium already overtaken by the wave, while remaining intact and identifiable in zones ahead of it (Fig. 4A, B). Coincident with their disappearance, a thick sulphated mucin layer appears on the outer surface of the collapsing epithelium (Fig. 4C, D). This suggests that globular cells rapidly discharge their contents outside the postlarva as the infolding wave passes through their zone, potentially providing a mucus-rich interface between epithelial surfaces being brought into apposition, and between the postlarva and the algal substrate and external environment. This mucus layer extends apically beyond the exocytosing globular cells, likely being spread by ciliary action of still functional ciliated columnar epithelial cells. This mass localised release of mucins within the morphogenetic wave contrasts with the distributed and measured release in swimming larvae (Fig. 2D, H and Supplementary Fig 1).

**Figure 4.**
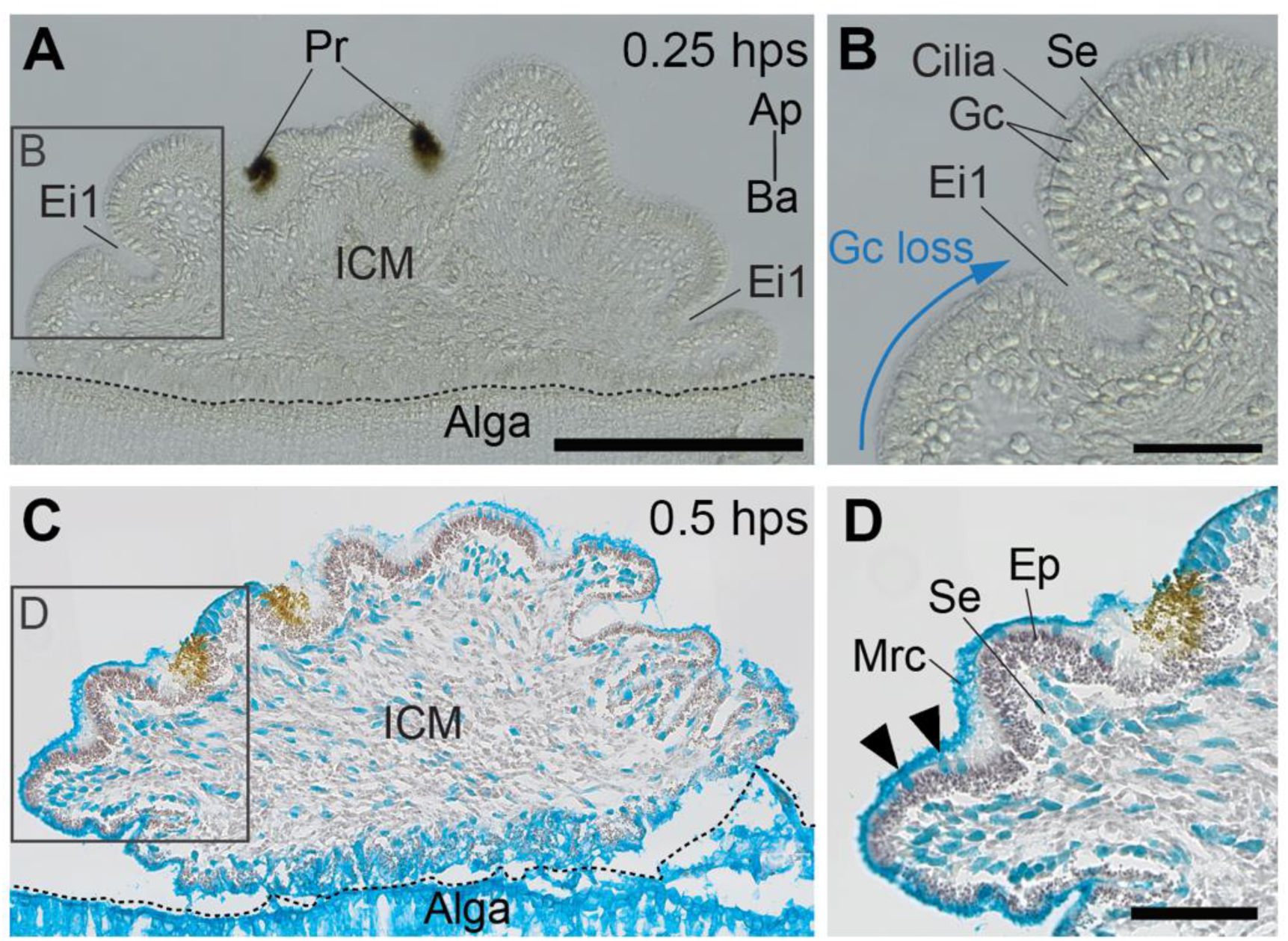
Rapid discharge of mucin from globular cells in concert with the morphogenetic wave. DIC micrographs of longitudinal sections of postlarvae settled on *A. fragilissima* (Alga); apical (Ap) – basal (Ba) axis shown. **(A, B)** Disappearance of globular cells (Gc) in 0.25 hps postlarva occurs as the first epithelial infolding (Ei1) advances, as marked by the blue arrow, while Gc above the infolding are still present. Subepithelial spherulous cells remain present in the subepithelial layer (Se). **(C, D)** 0.5 hps postlarvae stained for sulphated mucin with alcian blue (cyan) and hematoxylin (purple nuclei). Sulphated mucin is abundant on the surface of the postlarva and associated with the regions in contact with the alga and around resorbing cilia. Only a few globular cells containing mucin are still visible (black arrowheads). In contrast, subepithelial spherulous cells are still enriched in mucins. Scale bars: A, 200 µm; B, D, 50 µm.

### Targeted PCD eliminates larval-specific structures and culls larval epithelial cells

Low-level PCD in the vicinity of the pigment ring is already present in competent larvae prior to settlement (Fig. 2F), suggesting that these cells are primed for elimination before the metamorphic cue is received. Following settlement, PCD in the pigment ring increases progressively and by 6 hps most pigment ring cells appear to have undergone cell death (Fig. 5).

**Figure 5.**
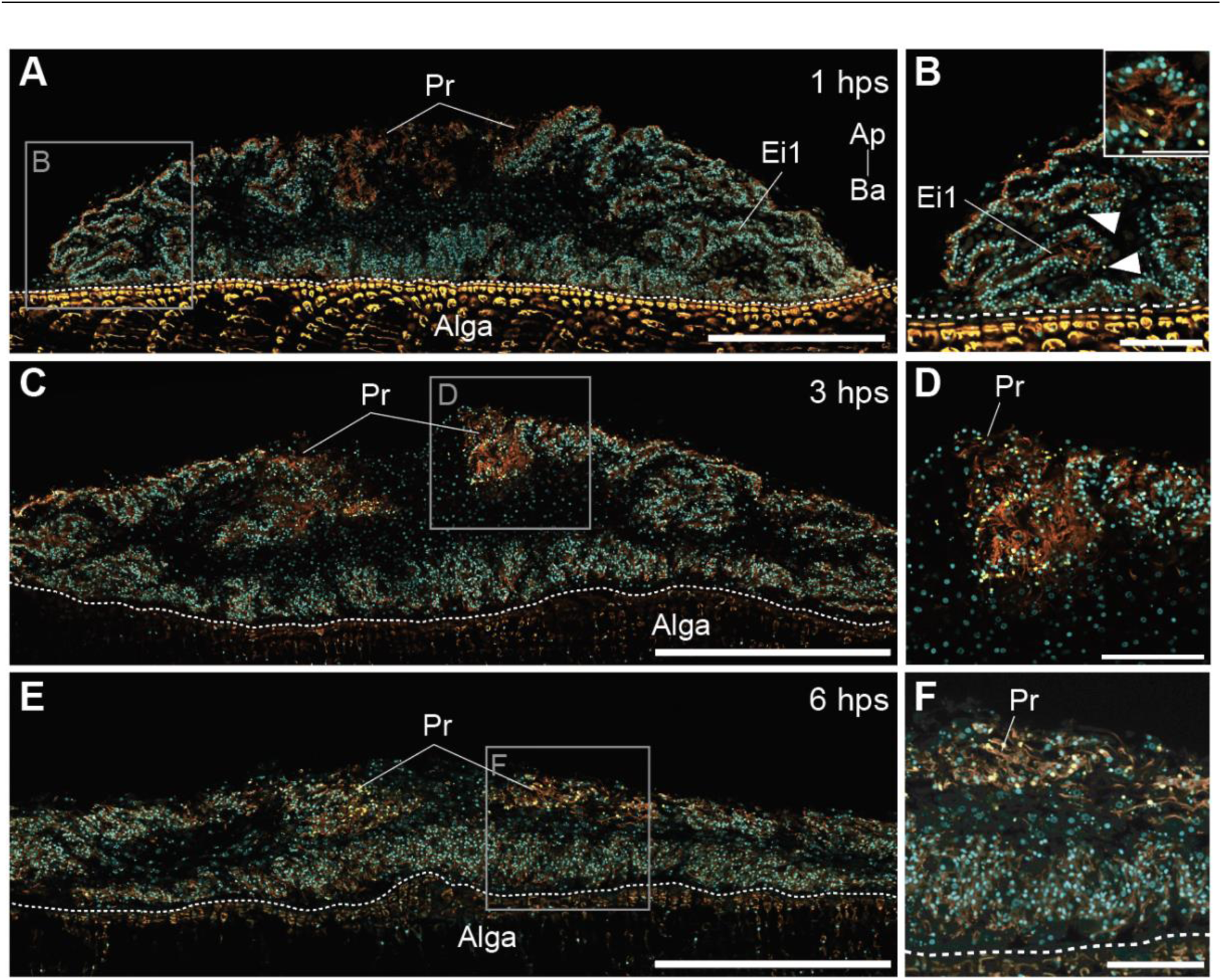
Programmed cell death of larval pigment ring and epithelial cells. Confocal micrographs of postlarvae as described in Fig. 3. **(A, B)** 1 hps postlarva with limited PCD occurring around the infolded epithelium and pigment ring. **(C, D)** Central region of a 3 hps postlarva with more extensive PCD near the receding pigment ring. **(E, F)** Central region of a 6 hps postlarva with PCD still occurring in the vicinity of the pigment ring. Scale bars: A, C, E, 200 µm; B, D, F, 50 µm.

The spatial confinement of PCD to the pigment ring and specific epithelial zones (Figs 3C, D and 5) is informative regarding induced changes at settlement and the initiation of metamorphosis. The pigment ring mediates larval phototaxis [40, 44] but is functionally dispensable in the sessile juvenile, making its elimination a clear case of targeted removal of a larval-specific structure. The pre-settlement priming of pigment ring cells for PCD is consistent with cell death fate being assigned during larval development, so that settlement then serves as a permissive trigger for a fate already determined.

### MET establishes a new postlarval epithelium from internal mesenchyme

Concurrent with the inward transitions described above, a complementary outward movement establishes a new postlarval surface epithelium (Fig. 6A-H). By 3 hps, non-ciliated spherulous and amoeboid cells, which are characterised by their high mucus content and originate from the larval ICM and subepithelial layers, begin to spread over the outer surface of the ingressing former larval ciliated epithelium (Fig. 6A-E). By 6 hps, these cells develop a filiform, squamous morphology, and form a layer covering the former larval epithelium and extending over the algal substrate (Fig. 6F-H).

**Figure 6.**
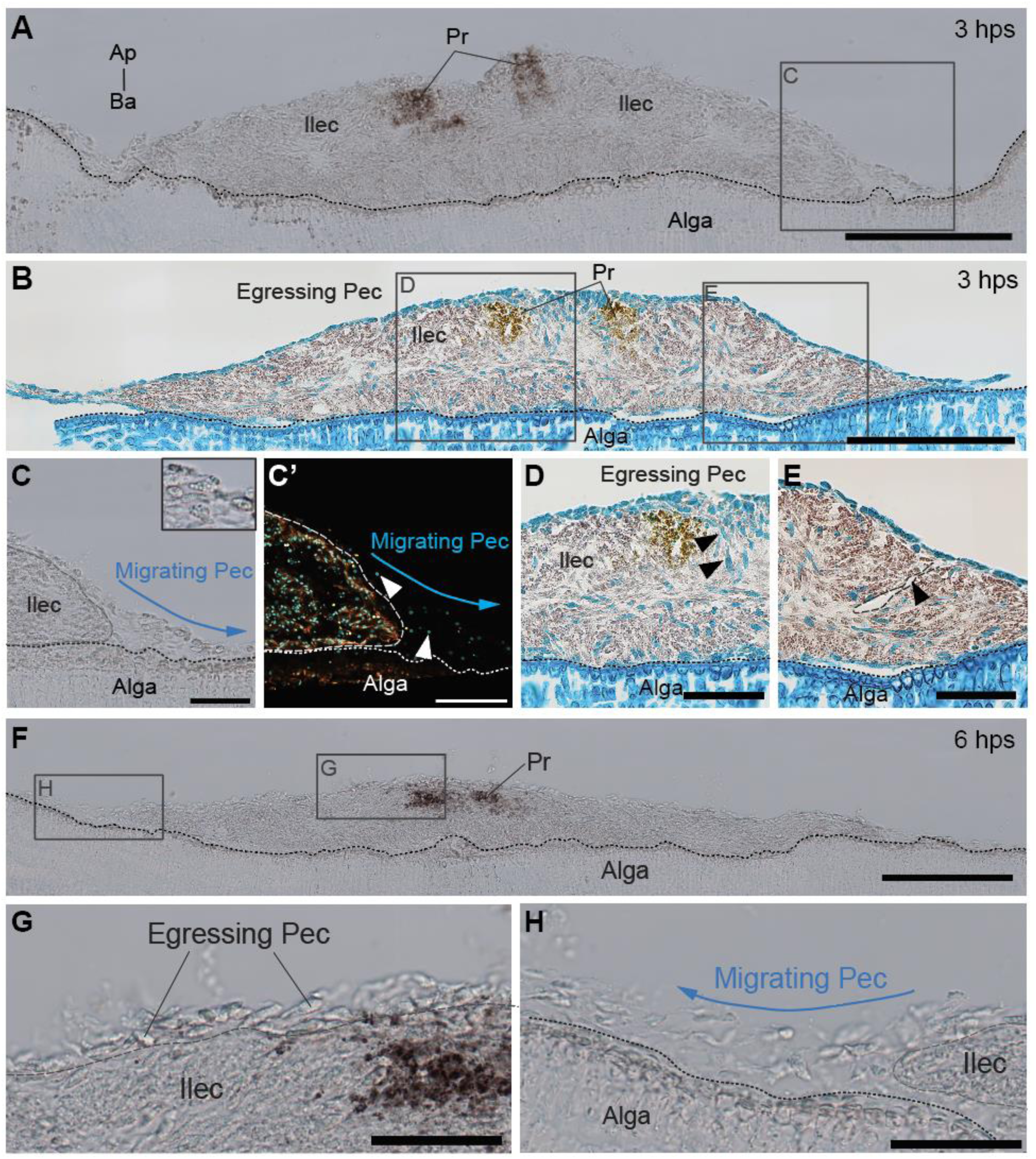
Emergence of the new postlarval epithelium. DIC and confocal micrographs of longitudinal sections of different postlarval stages. **(A, B)** 3 hps postlarvae with internal layer comprised of ingressing larval epithelial cells (Ilec) being covered by egressing mucin-rich spherulous-like cells (Pec), which are forming the postlarval epithelium. **(B)** Sulphated mucin stained with alcian blue (cyan) and nuclei with hematoxylin (purple). **(C, C’)** Outer edge of 3 hps postlarvae showing the spreading of postlarval epithelial cells (Pec) on the algal surface and recovering the Ilec (inside delimitations). Blue arrows, direction of cell migration; white arrowheads, Pec nuclei. **(D)** Mucin-rich spherulous-like cells concentrated under the former pigment ring (black arrowheads) and on the apical surface. **(E)** Mucin-rich epithelial cells spreading over the postlarval surface, enclosing former larval epithelial infoldings (arrowhead, dashed line). **(F-H)** 6 hps postlarva showing further accumulation and outward migration of postlarval epithelial cells. Scale bars: A, B, F, 200 µm; C-E, G, H, 50 µm.

This outward migration and surface colonisation is consistent with these mesenchymal cells acquiring epithelial identity and egressing to the postlarva exterior, potentially in the vicinity of the subsiding pigment ring (Fig. 6B, D). The resulting external layer is expected to become the juvenile exopinacoderm. The consequence of this MET, in conjunction with the inward movement of the former larval epithelium, is a wholesale exchange of cell layer identities within six hours of metamorphosis. The outer larval epithelium has been internalised and committed to choanocyte transdifferentiation, while internal larval mesenchyme cells have externalised and are establishing the new body surface. This inversion of layer identity is executed not by generating new cell populations but by repositioning and reprogramming existing ones. Interestingly, the thick mucus layer released at the start of metamorphosis is grown over by the egressing and migrating cells, which themselves are mucin-rich but do not appear to release mucus externally (Fig. 6B, D, E).

### Lineage tracing reveals the step-wise reprogramming of larval flask cells into archaeocytes

To track how a defined larval cell type is repurposed during metamorphosis, we labelled larval flask cells with the lipophilic dye CM-DiI and followed their fate from the competent larval stage through to 18 hps. Flask cells are pear-shaped, sensory-secretory cells enriched in the larval anterior half, bearing both apical and basal cytoplasmic projections and responding to settlement cues via intracellular calcium signalling (Figs 2E, I, 7A and Supplementary Fig. 1). We chose to track these cells specifically because these features make them strong candidates for initiating the metamorphic program [33].

By 0.5 hps, CM-DiI-labelled flask cells have become more evenly distributed across the postlarval body, likely a consequence of redistribution by the epithelial infoldings, and many have adopted an oval morphology with prominent intracellular vesicles (Fig. 7B), indicating early morphological change. By 3 hps, most labelled cells have an amoeboid morphology and have undergone EMT, localising within the subepithelium and ICM (Fig. 7C). A subset of labelled cells appears to have disintegrated at this stage (Fig. 7D), indicating that not all flask cells transdifferentiate, but some instead undergo cell death.

**Figure 7.**
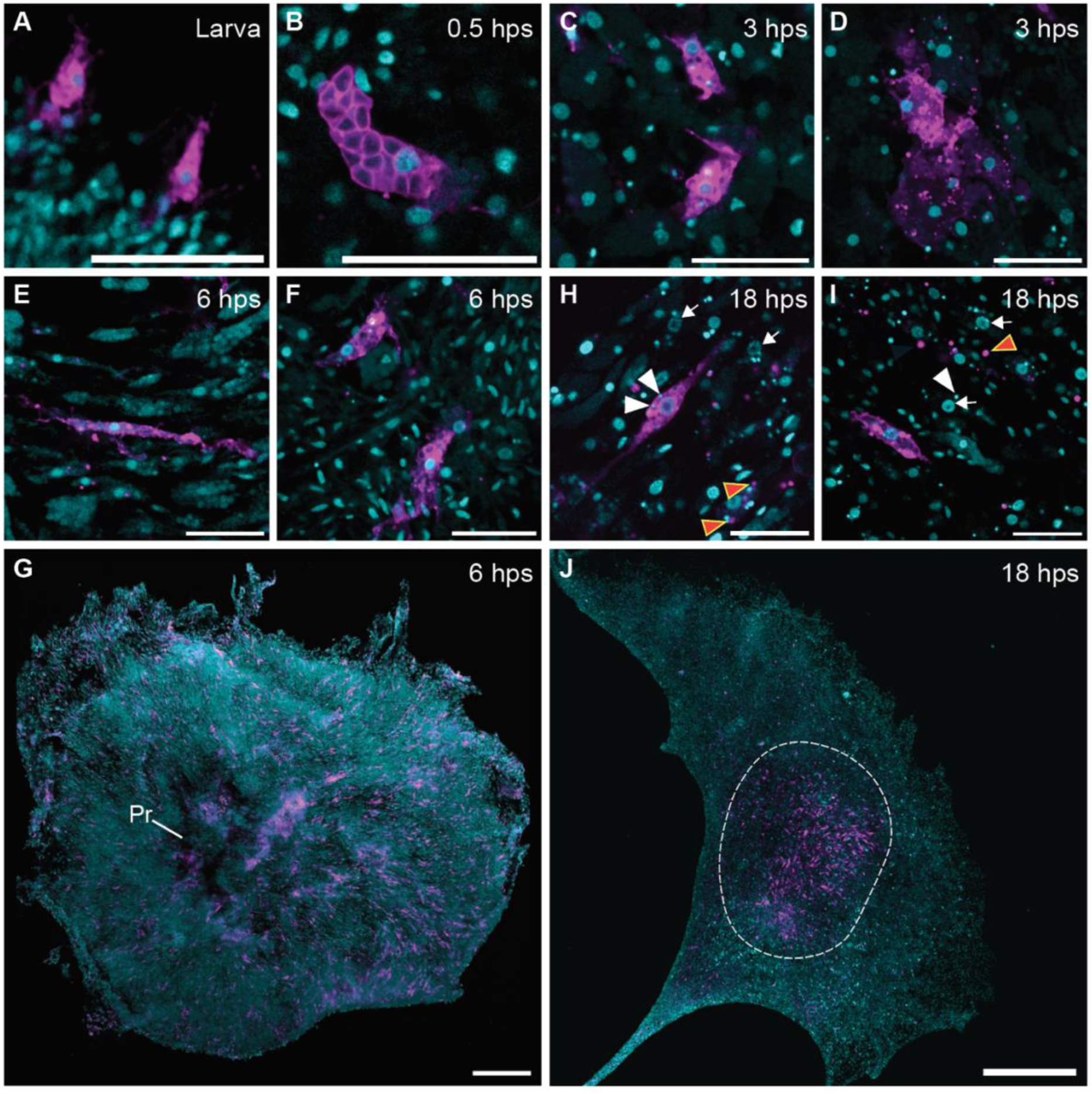
Transdifferentiation of larval flask cells during early metamorphosis. Confocal photomicrographs of whole-mount samples. Flask cells are labelled with CM-DiI (magenta) and all nuclei with Hoechst (cyan). **(A)** Flask cells are enriched in the epithelium of the anterior half of the competent swimming larva. **(B)** At 0.5 hps, many labelled flask cells adopt an oval shape with distinct vesicles and are more evenly distributed across the body. **(C, D)** At 3 hps, some CM-DiI-labelled cells appear to have undergone further morphological changes, resembling amoebocytes (C), while a few appear to be disintegrating (D). At this stage, flask cells are predomnantly internally localised consistent with having undergone EMT. **(E-G)** At 6 hps, CM-DiI-labelled cells are distributed throughout the apical region of the postlarva. **(E)** CM-DiI-labelled cells found at the leading edges of the postlarva on the alga are elongated and filiform with a central nucleus. **(F)** In contrast, CM-DiI-labelled cells in the apical centre of the postlarva are large and amoeboid. **(H-J)** At 18 hps, most CM-DiI-labelled cells are concentrated in the centre of the postlarva. **(H, I)** Most CM-DiI-labelled cells have transdifferentiated into archaeocytes, characterized by a large nucleus with a nucleolus. DiI-labelled and unlabelled archaeocytes (white arrows point to nuclei) appear to engulf DNA fragments (white arrowheads) and CM-DiI-labelled cytoplasmic fragments (orange arrowheads). Scale bars: A-F, H, I, 20 µm; G, 200 μm; J, 500 μm.

By 6 hps, most CM-DiI-labelled cells are concentrated internally in the apical region and are present in two morphologically distinct states: (i) elongated filiform cells at the spreading edge of the postlarval body, consistent with migratory mesenchymal cells contributing to the postlarva’s outward expansion; and (ii) large amoeboid cells within the postlarval interior, suggesting ongoing progression toward a larger cell type (Fig. 7E-G). By 18 hps, nearly all DiI-labelled cells have transformed into large archaeocytes with a prominent nucleus bearing a distinct nucleolus (Fig. 7H-I). Both labelled and unlabelled archaeocytes are concentrated centrally at this postlarval stage (Fig. 7H-J), indicating that flask cells are not the sole source of the archaeocyte pool. Indeed, numerous archaeocytes are present already in the larval ICM [14]. Many labelled and unlabelled archaeocytes at 18 hps contain small DNA and CM-DiI-labelled fragments within their cytoplasm, suggesting that phagocytosis of dying flask cell fragments occurs alongside successful transdifferentiation (Fig. 7H-I).

## Discussion

A central challenge in understanding metamorphosis is explaining not just which cellular processes are involved, but how they are orchestrated into a coherent program that reliably produces the juvenile body plan from the larval body plan. In marine invertebrates, this developmental transition is typically environmentally-induced and occurs rapidly. Our analysis of the first six hours of *Amphimedon queenslandica* metamorphosis offers the most temporally and spatially resolved picture of this orchestration in any sponge and reveals several organisational principles that are likely to have broader relevance to understanding metamorphic programs across animals.

### The wave of epithelial infoldings as a morphogenetic pacemaker

An important organisational feature of *A. queenslandica* early metamorphosis is the stereotypic basal-to-apical wave of epithelial infoldings that initiates within minutes of settlement and completes within one hour. In addition to collapsing the larval body, this wave appears to function as a spatial pacemaker that induces cellular transitions. Ciliary resorption of the columnar epithelial cells, increased exocytosis in globular epithelial cells, EMT of the flask epithelial cells, and early PCD each track with the position of the wavefront, occurring in zones already reached but not yet in zones ahead of it. This is consistent with that wave either directly triggering these events or propagating a signal that is the trigger. Although epithelial reorganisation at the start of metamorphosis occurs in other sponges [13, 20–22, 24–29, 45, 46], morphogenetic waves have not been previously document to our knowledge. Nonetheless, the coordinated epithelial movements that initiate the body plan transition in sponges have broad parallels in the diverse programmed tissue movements that initiate metamorphosis in parahoxozoans [1, 2, 6, 9, 47–54].

The initiation of this wave within minutes of the larva settling and attaching to the alga suggests that the primary inductive signal is from the alga itself or from a larval cell type immediately affected by an algal signal. Currently, the most likely initiating event is the anterior epithelial flask cells via calcium signalling [33, 41], although externally-facing cuboidal cells at the anterior pole also may be involved [35]. Flask cells possess apical and basal cytoplasmic projections resembling sensory neurons, positioning them to potentially receive the settlement cue first and transduce it to the surrounding epithelial cells, including globular cells [33]. The spatial coupling of globular cell exocytosis to the wavefront — globular cells vesicles disappear in zones already overtaken by the wave while remaining intact ahead of it — suggests that the wave itself, or a signal associated with it, triggers this event. Nitric oxide (NO), produced by globular cells, is a strong candidate for the propagating signal [41]. As a small, rapidly diffusing molecule, NO could activate epithelial remodelling in a spreading front from the point of initial larval attachment, potentially via flask cell activation. The progressive depletion of NO from globular cells could create a zone of high NO concentration that is associated with the wavefront, thus providing a self-sustaining propagation mechanism. Once the infoldings begin to bring previously distant epithelial surfaces into proximity, mechanical coupling through ciliary contacts could further coordinate the response [55]. The release of mucin may contribute to interactions between these newly apposed epithelial surfaces.

The wave’s directionality, which is maintained regardless of substrate geometry, suggests that the larva is spatially pre-patterned along its anteroposterior axis to ensure the direction of the morphogenetic program at the initiation of metamorphosis. This pre-patterning is likely maintained in the competent larva in a latent state, with settlement serving as the permissive trigger rather than the instructive inducer of spatial information. Pre-patterning of metamorphic programs in larvae is likely widespread in animals [1–3, 6–8, 54, 56–58]. The *A. queenslandica* system, with its tractable larval anatomy and inducible metamorphosis, offers a route to understanding how larval pre-patterning manifests at metamorphosis.

### Temporal induction of cell fate decisions

A crucial aspect of the morphogenetic program is that different cell populations undergo distinct state and fate changes in a defined temporal sequence rather than simultaneously, even when they are spatially intermingled. This temporal staggering is functionally important because if all cellular transitions were simultaneous, the postlarva would potentially risk catastrophic loss of tissue integrity. For instance, the epithelium cannot be simultaneously internalised and subjected to selective PCD without a coordinated handover. This morphogenetic sequencing appears to ensure that each cell state transition is contingent on, or at least compatible with, the state produced by preceding events. The infolding wave repositions the epithelium before transdifferentiation is underway, while subepithelial and ICM cells simultaneously undergo MET to colonise the surface being vacated by the internalising larval epithelium. These ordered transitions are a hallmark of a program rather than a set of independent responses.

The rapid induction of PCD in pigment ring cells is particularly informative in this respect. It suggests that these cells are already primed for PCD before the settlement cue is received and that temporal gating of some cell fate decisions is established during larval development, not at settlement. In these cases, settlement then acts as a coordinating signal that triggers the sequential execution of pre-assigned fates. The pre-assigned cell fates in the larva and their coordinated execution at metamorphosis may be a general principle of the pelagobenthic life cycle, allowing metamorphosis to proceed rapidly and reliably because cell specification has already occurred. It is also worth noting that the scale of PCD during early *A. queenslandica* metamorphosis appears quite limited. Cell death eliminates specific larval-specific structures but does not drive wholesale body remodelling, which is accomplished instead primarily by cell rearrangement, state change and layer exchange. PCD operates here as a targeted pruning mechanism running in parallel with the morphogenetic program, rather than as one of its primary drivers. This distinction may reflect a broader organisational principle in which the removal of larval structures is decoupled from the formation of juvenile ones.

### The archaeocyte as a reprogramming hub

The lineage tracing of flask cells reveals a defined sequence of state transitions: epithelial sensory cell to amoeboid migratory cell to archaeocyte stem cell. This sequence constitutes a directed reprogramming pathway in which the amoeboid intermediate functions as a morphological, and presumably molecular, gateway between the larval epithelial identity and the pluripotent archaeocyte identity. The archaeocyte, once formed, can differentiate into diverse juvenile cell types including choanocytes and pinacocytes [17, 19, 59–62], making the flask cell-to-archaeocyte transition a cellular bridge between the larval and juvenile body plans. Whether archaeocytes derived from flask cells preferentially adopt specific juvenile cell types or are equivalent to archaeocytes derived from other larval cell types, remains to be determined. The observation that some flask cells die rather than transdifferentiate further underscores that while the population-level program is stereotypic individual cell outcomes are not deterministic. Archaeocyte phagocytosis of these dying cell fragments may serve a dual role, clearing cellular debris while simultaneously transferring molecular contents between cells.

This phenomenon is reminiscent of dedifferentiation-redifferentiation sequences in bilaterian metamorphosis and regeneration, where cells transiently pass through a less differentiated state to a new identity [63–65]. In *A. queenslandica*, the archaeocyte may serve as the universal convergence point for multiple larval cell types, including flask cells and some columnar epithelial cells, before they diverge again into the specific juvenile cell types. This would make the archaeocyte not just a stem cell but a reprogramming hub of metamorphosis where multiple larval cell states converge upon before juvenile cell type diversity emerges. This is consistent with archaeocyte pluripotency as a general principle of sponge cell biology [21, 30, 66, 67]. In addition to transdifferentiation, the postlarva inherits a large population of archaeocytes located in the larval ICM, lending support to the premise that archaeocyte stem cells, whether a cell state intermediate or not, are critical to the generation of juvenile cell types and body plan. Whether archaeocytes derived from different larval cell types are molecularly equivalent to these larval-derived archaeocytes or retain lineage-specific information that biases their subsequent differentiation, is an important open question. Cell-type-resolved transcriptomics of the archaeocyte population during metamorphosis may shed light on this question.

### Integrating conserved cellular processes into a body plan switch

Each cellular process observed during *A. queenslandica* early metamorphosis – epithelial infoldings, EMT, MET, transdifferentiation, PCD, coordinated secretion – individually is a conserved feature of animal development. EMT and MET are fundamental to gastrulation, organogenesis and wound healing across the Parahoxozoa [68–70]. Transdifferentiation occurs in cnidarian and bilaterian metamorphosis [10, 11, 64, 71], as well as in sponge regeneration [60, 61, 67, 72]. PCD at metamorphosis is widespread across marine invertebrates [73–77]. The *A. queenslandica* case presented here confirms the existence of these processes in sponge metamorphosis, but also goes much further by revealing the organisational logic by which they are integrated into a coherent body plan transition.

Three features of this integration stand out. First, the wave of epithelial infoldings provides a spatiotemporal framework for the program, ensuring that different cellular transitions occur in the correct anatomical zones and in the correct sequence relative to one another. Second, some cell fate decisions appear to be pre-assigned in the larva and are triggered in a defined temporal order at settlement, preventing the chaos that could result from simultaneous execution of conflicting cellular transitions. Third, the convergence of multiple larval cell types on a common pluripotent archaeocyte intermediately before further differentiation provides a flexible mechanism for generating diverse juvenile cell types from larval cell types without requiring direct transdifferentiation of each larval type into its juvenile counterpart (e.g., larval ciliated columnar cells into juvenile choanocytes).

Whether these three organisational principles are general features of animal metamorphosis, or specific to the sponge lineage, is an important question for comparative developmental biology. The pelagobenthic life cycle is ancient and widespread, and the pressure to execute the larva-to-juvenile transition rapidly and reliably is shared across the lineages that employ it [2]. It would not be surprising if different animal phyla have converged on similar organisational solutions, even if cell types, signals and molecular mechanisms differ. The cellular framework established here for *A. queenslandica* metamorphosis provides a means to understand organisational features in other animals, allowing for the identification and understanding of shared principles underlying metamorphosis.

## Methods

### Sample collection

Adult *Amphimedon queenslandica* were collected from Heron Island Reef and translocated to the aquarium facility at the University of Queensland within 2 days of being collected [78, 79]. Larval release was triggered by a gradual rise in seawater temperature from 25.5°C to 27.5°C for 2 h, starting at 11:00 a.m. Swimming larvae were collected into a beaker of filtered seawater (FSW) and maintained at ambient temperature (25.5°C) and light for 6 h. Most of swimming larvae became competent to settle and metamorphose at the time of sunset, 7:00 pm [32, 34]. Competent larvae were transferred to 6-well plates (CLS3516 Corning Costar) containing 5 mL of FSW and four small (∼1 cm in length) Y-shaped pieces of brush-cleaned live coralline algae *Amphiroa fragilissima* per well. *A. fragilissima* was also collected from Heron Island Reef. The prepared plates were then placed in the dark to induce settlement. One hour later, unsettled larvae were removed, and the settled postlarvae were raised to the age of interest. At this stage, the newly settled postlarvae were between 0.5 − 1 hps; these early postlarvae have different external morphologies allowing them to be accurately staged.

### Video recording of the initiation of metamorphosis

Competent larvae were induced to settle and metamorphose on *A. fragilissima*. After 30 min in the dark, newly settled larvae were selected to time-lapse video record the start of metamorphosis. Images of *A. queenslandica* postlarvae were taken with a Nikon SMZ25 microscope using the program Nikon Elements and approximately 6X zoom. For Supplementary video 1, images were taken every 3 min, starting from approximately 10 min after the initiation of settlement through to approximately 2 hps. Each time point consisted of a z-stack with 14 images at an interval of 25 µm, which were then stacked into a single image using the focus stacking program Zerene Stacker with the P Max algorithm ^48^. These were cropped, brightened and denoised, then unsharp masked using Adobe Photoshop. To generate the timelapse movie, the frames were compiled into an avi using ImageJ and converted to mp4 (or mkv) using AnyVideoConverter (https://www.any-video-converter.com/en8/for_video_free/). Topaz Video Enhance AI (Topaz Labs) was used to achieve AI-mediated temporal interpolation to generate 960 frames for smoother presentation at 24 frames per second (fps); the 1 fps raw data movie is available upon request. The movie is shown at 180x normal speed. For Supplementary videos 2-4, single images (no z-stack) were taken every 1 min for 1 – 1.5 h, starting from approximately 10 min after the initiation of settlement through to approximately 1 hps. Images were cropped, brightened and compiled into an avi using ImageJ, and were converted to mp4 using AnyVideoConverter; the movies are shown at 245x normal speed.

### Fixation and cryosectioning

Competent larvae and postlarvae on *A. fragilissima* were fixed in 4% paraformaldehyde (PFA) and 0.05% glutaraldehyde in modified 3-(N-morpholino) propane sulfonic acid buffer (MOPS pH 7.5: 0.1 M MOPS; 0.5 M NaCl; and 2 mM MgSO4) at room temperature (RT) on a slow nutator for 1 h, then dehydrated in 70% ethanol and stored at −20°C until use [78, 80].

After gradual rehydration from 70% ethanol to distilled water, postlarvae settled on algae were incubated in 0.25 M egtazic acid (EGTA) pH 8 for 24 – 48 h at RT to decalcify the algae. Then, an equal volume of 30% sucrose was added to the existing solution and gently mixed by pipetting for 1 min. The solution was then completely removed and replaced with fresh 30% sucrose, and samples were placed on a slow nutator for 15 min. Samples were transferred into a cryo-mould and quickly washed three times with 9% gelatine + 10% sucrose. The latter was added again, and the specimens were rapidly oriented before embedding. The cryo-mould was then covered with a lid, and a prechilled −80°C block was placed on top of the lid. Once the gelatine had set, the cryoblock was stored at −80°C until use. The cryoblock was sectioned using a Cryostat Microm HM560, and the sections were mounted on adhesive-coated Uber slides (instrumeC).

### Immunofluorescence and TUNEL

Cryosections were incubated in pre-warmed MOPS (37°C) for 1 min to remove the gelatine. Samples were permeabilised through three 5 min washes in phosphate-buffered saline (PBS) supplemented with 0.1% Tween® 20 (PBST), and blocked in 1% bovine serum albumin (BSA) in PBST for 30 min at RT. To label cilia, sections were first incubated with a mouse anti-acetyl-α-tubulin primary antibody (MABT868 Sigma-Aldrich) diluted 1:200 in 1% BSA PBST for 1 h at RT, washed three times for 5 min each in 1% BSA PBST, and then incubated with a goat anti-mouse secondary antibody conjugated to AlexaFluor 647 (A32728 Invitrogen) diluted 1:500 in 1% BSA PBST for 1 h at RT in the dark [17, 18]. After three 5-min washes in PBS, programmed cell death was assessed using TUNEL assay (TMR red 12156792910 Roche). Sections were incubated in the TUNEL reaction mixture (enzyme solution and label solution diluted 1:10) for 1 h at RT and washed three times for 5 min each in PBS. Nuclei were counterstained with Hoechst 33342 (1:1000) for 5 min, and samples were mounted in Prolong Glass Antifade Mountant (P36980 Invitrogen). Images were acquired using a Zeiss LSM 900 confocal microscope and processed using ImageJ.

### Transmission electron microscopy

Postlarvae at ∼1 hps, settled on *A. fragilissima*, were fixed as described above and were imaged using a NeoScope JCM-5000 Jeol benchtop scanning electron microscope.

### Alcian blue staining

Competent larvae and postlarvae were processed for sulphated mucin detection using alcian blue pH 1. Specimens were first fixed and stored in 70% ethanol at −20°C,then rehydrated in distilled water and incubated overnight at RT in 0.25 M citric acid to decalcify the algae on which they had settled; EGTA was not used here, as it interferes with alcian blue binding. Samples were incubated in 30% sucrose for 30 min, then embedded in a 1:1 mixture of OCT (Optimal Cutting Temperature) embedding medium and 30% sucrose. Cryoblocks were sectioned using a Cryostat Microm HM560, and sections were mounted on adhesive-coated Uber slides (InstrumeC), which have been further coated to improve section adherence as follows: coated with 2% (3-Aminopropyl) triethoxysilane (APTES; 440140 Sigma-Aldrich) for 10 min, air-dried, then coated with 0.05% glutaraldehyde for 2 min and air-dried. Cryosections were stored at −20°C until use.

Cryosections were rehydrated in distilled water for 5 min, immersed in 0.1 N HCl for 3 min, then incubated overnight at RT in 2% alcian blue (05500 Sigma-Aldrich), dissolved in 0.1 N HCl and filtered through a 0.22 µm filter). Sections were rinsed briefly in 10% methanol in 0.1 N HCl, followed by a 5 min wash in distilled water. Nuclei were counterstained with Weigert’s hematoxylin, and slides were mounted in Depex mounting medium (3612540 BDH). Images were acquired using an Olympus microscope in white light and processed using ImageJ software.

### Flask cell lineage tracing with CM-DiI

Live competent larvae were incubated in 10 µM CM-DiI dye (C7000 Invitrogen) in FSW for 2 h in the dark to selectively label flask cells [17, 18]. They were then washed three times for 5 min each in FSW, after which some of these larvae were fixed in 4% PFA and 0.2% glutaraldehyde in MOPS at RT on a slow nutator for 1 h, while the remaining larvae were induced to settle and metamorphose on *A. fragilissima*. Postlarvae were fixed at 0.5, 1, 3, 6 and 18 hps, as described above for the competent larvae. For the 18 hps postlarval stage, newly settled postlarvae (∼1 hps) were peeled off the algae, transferred onto a coverslip and fixed 18 h later. After fixation, samples were washed three times in MOPS for 5 min each, nuclei-counterstained with Hoechst 33342 (1:1000) and whole-mounted in Prolong Glass Antifade Mountant (P36980 Invitrogen); 0.5, 1, 3 and 6 hps postlarvae were peeled off the algae before mounting. Images were acquired within hours to two days after mounting using a Zeiss LSM 900 confocal microscope and processed using ImageJ.

### Phalloidin and cresyl violet staining

Live competent larvae were incubated in 1:500 cresyl violet (C1791 Sigma-Aldrich, solution stock is 1 mg/ml in distilled water) for 2 min in the dark and were then briefly washed in FSW. Samples were subsequently kept in the dark. Samples were fixed in 4% PFA, 0.05% glutaraldehyde in MOPS for 1 h at RT, and washed 3 times for 5 min in MOPS. After this, samples were incubated in 1:100 phalloidin (A12379 Invitrogen, dissolved in methanol) for 30 min, washed 3 times for 5 min in MOPS, and prepared for cryosectioning as described above. Cryosections were incubated in pre-warmed MOPS (37°C) supplemented with 1:50000 Hoechst 33342 for 5 min, then mounted in MOPS and imaged immediately using a Zeiss LSM 900 confocal microscope. The resulting images were processed using ImageJ.

## Acknowledgements

We acknowledge the support of the Heron Island Research Station staff in the collection and maintenance of biological materials, the Queensland Brain Institute for assistance with microscopy and thank Leïla Kopplin for technical assistance.

## Author information

OB designed the project, performed most laboratory and microscopy analyses and wrote the paper. ZP undertook laboratory and microscopy analyses of alcian blue, cresyl violet and phalloidin staining, and commented upon the paper. ST contributed to the project design and commented upon the paper. SMD designed, coordinated and supervised the project and wrote the manuscript. BMD designed, coordinated and supervised the project and wrote the manuscript. All authors approved the submitted version of the paper.

## Supplementary information

**Supplementary Fig. 1.**
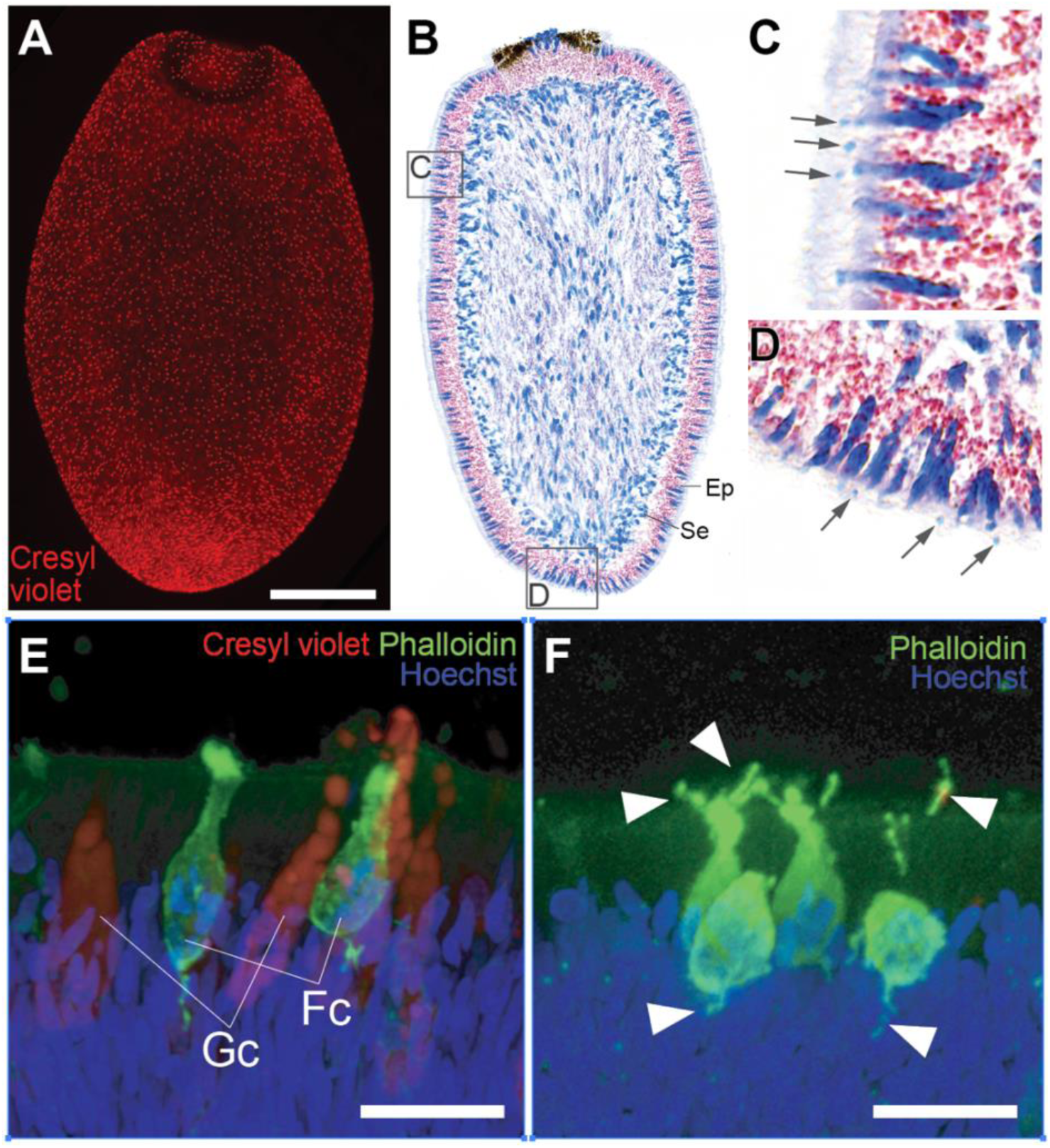
Larval globular and flask cells. **(A)** Whole mount fluorescence micrograph of competent larva stained with cresyl violet (red) that labels globular cells red. These are distributed evenly over the larval surface. **(B)** DIC micrograph of longitudinal section of competent larva stained with alcian blue (cyan) and hematoxylin (purple) showing the distribution of mucin-rich globular cells within the epithelial layer (Ep) and spherulous cells in the subepithelial layer (Se). **(C, D)** Magnifications of regions boxed in B showing the release of mucin globules outside the larva (arrows) that appear to coat the larval cilia. **(E, F)** Confocal micrographs of longitudinal sections of competent larvae stained with cresyl violet (red), phalloidin (green) and Hoechst 33342 (blue) showing the relationship of flask cells (Fc; green), globular (Gc; red) to each other and the columnar epithelium (blue nuclei and light green background). Arrowheads, some flask cell apical and basal processes. Scale bars: A, 100 µm; E, F, 10 µm.

**Supplementary videos 1-4.** Four time lapse videos of the first hours of metamorphosis. In all cases, a *A. queeslandica* larva has attached to and settled upon a fragment of the coralline alga, *A. fragilissima* (see Methods for details).

## References

1. Jägersten G. Evolution of the metazoan life cycle: a comprehensive theory. 1972.

2. Hadfield MG. Why and how marine-invertebrate larvae metamorphose so fast. Seminars in Cell & Developmental Biology. 2000;11:437–43. 10.1006/scdb.2000.0197.

3. Heyland A, Moroz LL. Signaling mechanisms underlying metamorphic transitions in animals. Integrative and Comparative Biology. 2006;46:743–59. 10.1093/icb/icl023.

4. Ereskovsky AV, Renard E, Borchiellini C. Cellular and molecular processes leading to embryo formation in sponges: evidences for high conservation of processes throughout animal evolution. Dev Genes Evol. 2013;223:5–22. 10.1007/s00427-012-0399-3.

5. Lv Z, Lu Q, Dong B. Morphogenesis: a focus on marine invertebrates. Mar Life Sci Technol. 2019;1:28–40. 10.1007/s42995-019-00016-z.

6. Fuchs J, Martindale MQ, Hejnol A. Gene expression in bryozoan larvae suggest a fundamental importance of pre-patterned blastemic cells in the bryozoan life-cycle. EvoDevo. 2011;2:13. 10.1186/2041-9139-2-13.

7. Formery L, Lowe CJ. Integrating Complex Life Cycles in Comparative Developmental Biology. Annual Review of Genetics. 2023;57 Volume 57, 2023:321–39. 10.1146/annurev-genet-071719-020641.

8. Pieplow C, Furze A, Gregory P, Oulhen N, Wessel GM. Sex specific gene expression is present prior to metamorphosis in the sea urchin. Developmental Biology. 2025;517:217–33. 10.1016/j.ydbio.2024.10.003.

9. Wray GA. Gene expression during echinoderm metamorphosis. Zygote. 1999;8:S48–9. 10.1017/S0967199400130230.

10. Krasovec G, Frank U. Apoptosis-dependent head development during metamorphosis of the cnidarian *Hydractinia symbiolongicarpus*. Developmental Biology. 2024;516:148–57. 10.1016/j.ydbio.2024.08.010.

11. Yuan D, Nakanishi N, Jacobs DK, Hartenstein V. Embryonic development and metamorphosis of the scyphozoan *Aurelia*. Dev Genes Evol. 2008;218:525–39. 10.1007/s00427-008-0254-8.

12. Grasso LC, Negri AP, Fôret S, Saint R, Hayward DC, Miller DJ, et al. The biology of coral metamorphosis: Molecular responses of larvae to inducers of settlement and metamorphosis. Developmental Biology. 2011;353:411–9. 10.1016/j.ydbio.2011.02.010.

13. Ereskovsky AV, Konyukov PYu, Tokina DB. Morphogenesis accompanying larval metamorphosis in *Plakina trilopha* (Porifera, Homoscleromorpha). Zoomorphology. 2010;129:21–31. 10.1007/s00435-009-0097-5.

14. Leys SP, Degnan BM. Embryogenesis and metamorphosis in a haplosclerid demosponge: gastrulation and transdifferentiation of larval ciliated cells to choanocytes. Invertebrate Biology. 2002;121:171–89. 10.1111/j.1744-7410.2002.tb00058.x.

15. Amano S, Hori I. Transdifferentiation of Larval Flagellated Cells to Choanocytes in the Metamorphosis of the Demosponge *Haliclona permollis*. The Biological Bulletin. 1996;190:161–72. 10.2307/1542536.

16. Amano S, Hori I. Metamorphosis of Coeloblastula Performed by Multipotential Larval Flagellated Cells in the Calcareous Sponge *Leucosolenia laxa*. The Biological Bulletin. 2001;200:20–32. 10.2307/1543082.

17. Nakanishi N, Sogabe S, Degnan BM. Evolutionary origin of gastrulation: insights from sponge development. BMC Biology. 2014;12:26. 10.1186/1741-7007-12-26.

18. Sogabe S, Nakanishi N, Degnan BM. The ontogeny of choanocyte chambers during metamorphosis in the demosponge *Amphimedon queenslandica*. EvoDevo. 2016;7:6. 10.1186/s13227-016-0042-x.

19. Sogabe S, Hatleberg WL, Kocot KM, Say TE, Stoupin D, Roper KE, et al. Pluripotency and the origin of animal multicellularity. Nature. 2019;570:519–22. 10.1038/s41586-019-1290-4.

20. Efremova S. Once more on the position among Metazoa—gastrulation and germinal layers of sponges. Berliner Geowiss Abh. 1997;20:7–15.

21. Ereskovsky A. The Comparative Embryology of Sponges. Dordrecht: Springer Netherlands; 2010. 10.1007/978-90-481-8575-7.

22. Leys SP, Ereskovsky AV. Embryogenesis and larval differentiation in sponges. Can J Zool. 2006;84:262–87. 10.1139/z05-170.

23. Borisenko I, Podgornaya OI, Ereskovsky AV. Chapter Twelve - From traveler to homebody: Which signaling mechanisms sponge larvae use to become adult sponges? In: Donev R, editor. Advances in Protein Chemistry and Structural Biology. Academic Press; 2019. p. 421–49. 10.1016/bs.apcsb.2019.02.002.

24. Maldonado M. The ecology of the sponge larva. Canadian Journal of Zoology. 2006;84.

25. Delage Y. Embryogenèse des éponges siliceuses. Archs Zool Exp Gén. 1892;10:345–498.

26. Gonobobleva E, Ereskovsky A. Metamorphosis of the larva of *Halisarca dujardini* (Demospongiae, Halisarcida). Boll Mus Ist Biol Univ Genova. 2004;74:101–15.

27. Ereskovsky AV, Konjukov P, Willenz P. Experimental metamorphosis of *Halisarca dujardini* larvae (Demospongiae, Halisarcida): Evidence of flagellated cell totipotentiality. Journal of Morphology. 2007;268:529–36. 10.1002/jmor.10481.

28. Mukhina YI, Kumeiko VV, Podgornaya OI, Efremova SM. The fate of larval flagellated cells during metamorphosis of the sponge *Halisarca dujardini*. Int J Dev Biol. 2006;50:533–41. 10.1387/ijdb.052123ym.

29. Ereskovsky AV, Tokina DB, Bézac C, Boury-Esnault N. Metamorphosis of cinctoblastula larvae (Homoscleromorpha, porifera). Journal of Morphology. 2007;268:518–28. 10.1002/jmor.10506.

30. Simpson TL. The Cell Biology of Sponges. Springer Science & Business Media; 1984.

31. Maldonaldo M, Wahub MAA. Phylum Porifera. In: Atlas of Marine Invertebrate Larvae. Academic Press; 2025. p. 113–48. 10.1016/B978-0-08-102871-1.00002-6.

32. Degnan SM, Degnan BM. The initiation of metamorphosis as an ancient polyphenic trait and its role in metazoan life-cycle evolution. Philosophical Transactions of the Royal Society B: Biological Sciences. 2010;365:641–51. 10.1098/rstb.2009.0248.

33. Nakanishi N, Stoupin D, Degnan SM, Degnan BM. Sensory Flask Cells in Sponge Larvae Regulate Metamorphosis via Calcium Signaling. Integrative and Comparative Biology. 2015;55:1018–27. 10.1093/icb/icv014.

34. Say TE, Degnan SM. Molecular and behavioural evidence that interdependent photo - and chemosensory systems regulate larval settlement in a marine sponge. Molecular Ecology. 2020;29:247–61. 10.1111/mec.15318.

35. Degnan BM, Adamska M, Richards GS, Larroux C, Leininger S, Bergum B, et al. Porifera. In: Wanninger A, editor. Evolutionary Developmental Biology of Invertebrates 1: Introduction, Non-Bilateria, Acoelomorpha, Xenoturbellida, Chaetognatha. Vienna: Springer; 2015. p. 65–106. 10.1007/978-3-7091-1862-7_4.

36. Sebé-Pedrós A, Chomsky E, Pang K, Lara-Astiaso D, Gaiti F, Mukamel Z, et al. Early metazoan cell type diversity and the evolution of multicellular gene regulation. Nat Ecol Evol. 2018;2:1176–88. 10.1038/s41559-018-0575-6.

37. Yuan H, Blard O, Pujic Z, Degnan BM, Degnan SM. Light-entrained chromatin priming poises rapid metamorphosis in a marine sponge. 2026;:2026.01.28.702437. 10.64898/2026.01.28.702437.

38. Richards GS, Simionato E, Perron M, Adamska M, Vervoort M, Degnan BM. Sponge Genes Provide New Insight into the Evolutionary Origin of the Neurogenic Circuit. Current Biology. 2008;18:1156–61. 10.1016/j.cub.2008.06.074.

39. Richards GS, Degnan BM. The expression of Delta ligands in the sponge *Amphimedon queenslandica* suggests an ancient role for Notch signaling in metazoan development. EvoDevo. 2012;3:15. 10.1186/2041-9139-3-15.

40. Leys SP, Degnan BM. Cytological Basis of Photoresponsive Behavior in a Sponge Larva. The Biological Bulletin. 2001;201:323–38. 10.2307/1543611.

41. Ueda N, Richards GS, Degnan BM, Kranz A, Adamska M, Croll RP, et al. An ancient role for nitric oxide in regulating the animal pelagobenthic life cycle: evidence from a marine sponge. Sci Rep. 2016;6:37546. 10.1038/srep37546.

42. Martin GG. Ciliary gliding in lower invertebrates. Zoomorphologie. 1978;91:249–61. 10.1007/BF00999814.

43. Kuek LE, Lee RJ. First contact: the role of respiratory cilia in host-pathogen interactions in the airways. American Journal of Physiology-Lung Cellular and Molecular Physiology. 2020;319:L603–19. 10.1152/ajplung.00283.2020.

44. Wong E, Anggono V, Williams SR, Degnan SM, Degnan BM. Phototransduction in a marine sponge provides insights into the origin of animal vision. iScience. 2022;25:104436. 10.1016/j.isci.2022.104436.

45. Leys SP, Eerkes-Medrano D. Gastrulation in Calcareous Sponges: In Search of Haeckel’s Gastraea1. Integrative and Comparative Biology. 2005;45:342–51. 10.1093/icb/45.2.342.

46. Eerkes-Medrano DI, Leys SP. Ultrastructure and embryonic development of a syconoid calcareous sponge. Invertebrate Biology. 2006;125:177–94. 10.1111/j.1744-7410.2006.00051.x.

47. Weis VM, Buss LW. Ultrastructure of metamorphosis in Hydractinia echinata. 1987.

48. Kraus YA. Cnidarian Larvae: True Planulae, Other-Than-Planulae, and Planulae That Don’t Look Like Planulae. Russ J Dev Biol. 2023;54:S23–61. 10.1134/S1062360423070044.

49. Cameron RA, Hinegardner RT. Early events in sea urchin metamorphosis, description and analysis. Journal of Morphology. 1978;157:21–31. 10.1002/jmor.1051570103.

50. Reed CG, Cloney RA. The settlement and metamorphosis of the marine bryozoan *Bowerbankia gracilis* (Ctenostomata: Vesicularioidea). Zoomorphology. 1982;101:103–32. 10.1007/BF00312018.

51. Reed CG, Woollacott RM. Mechanisms of rapid morphogenetic movements in the metamorphosis of the bryozoan *Bugula neritina* (cheilostomata, cellularioidea). I. Attachment to the substratum. Journal of Morphology. 1982;172:335–48. 10.1002/jmor.1051720308.

52. Stricker SA. Metamorphosis of the marine bryozoan *Membranipora membranacea*: An ultrastructural study of rapid morphogenetic movements. Journal of Morphology. 1988;196:53–72. 10.1002/jmor.1051960106.

53. Bonar DB. Molluscan Metamorphosis: A Study in Tissue Transformation. American Zoologist. 1976;16:573–91.

54. Gonzalez P, Jiang JZ, Lowe CJ. The development and metamorphosis of the indirect developing acorn worm *Schizocardium californicum* (Enteropneusta: Spengelidae). Front Zool. 2018;15:26. 10.1186/s12983-018-0270-0.

55. Bloodgood RA. Sensory reception is an attribute of both primary cilia and motile cilia. Journal of Cell Science. 2010;123:505–9. 10.1242/jcs.066308.

56. Degnan BM, Groppe JC, Morse DE. Chymotrypsin mRNA expression in digestive gland amoebocytes: cell specification occurs prior to metamorphosis and gut morphogenesis in the gastropod, Haliotis rufescens. Roux’s Arch Dev Biol. 1995;205:97–101. 10.1007/BF00188848.

57. Degnan BM, Morse DE. Developmental and Morphogenetic Gene Regulation in *Haliotis rufescens* Larvae at Metamorphosis. American Zoologist. 1995;35:391–8.

58. Maslakova SA. Development to metamorphosis of the nemertean pilidium larva. Frontiers in Zoology. 2010;7:30. 10.1186/1742-9994-7-30.

59. Funayama N, Nakatsukasa M, Mohri K, Masuda Y, Agata K. *Piwi* expression in archeocytes and choanocytes in demosponges: insights into the stem cell system in demosponges. Evolution & Development. 2010;12:275–87. 10.1111/j.1525-142X.2010.00413.x.

60. Borisenko IE, Adamska M, Tokina DB, Ereskovsky AV. Transdifferentiation is a driving force of regeneration in *Halisarca dujardini* (Demospongiae, Porifera). PeerJ. 2015;3:e1211. 10.7717/peerj.1211.

61. Funayama N. The cellular and molecular bases of the sponge stem cell systems underlying reproduction, homeostasis and regeneration. Int J Dev Biol. 2018;62:513–25. 10.1387/ijdb.180016nf.

62. Funayama N. The stem cell system in demosponges: suggested involvement of two types of cells: archeocytes (active stem cells) and choanocytes (food-entrapping flagellated cells). Dev Genes Evol. 2013;223:23–38. 10.1007/s00427-012-0417-5.

63. Jopling C, Boue S, Belmonte JCI. Dedifferentiation, transdifferentiation and reprogramming: three routes to regeneration. Nat Rev Mol Cell Biol. 2011;12:79–89. 10.1038/nrm3043.

64. Hirano T, Nishida H. Developmental Fates of Larval Tissues after Metamorphosis in Ascidian *Halocynthia roretzi*. Developmental Biology. 1997;192:199–210. 10.1006/dbio.1997.8772.

65. Tiozzo S, Copley RR. Reconsidering regeneration in metazoans: an evo-devo approach. Front Ecol Evol. 2015;3. 10.3389/fevo.2015.00067.

66. Ereskovsky A, Melnikov NP, Lavrov A. Archaeocytes in sponges: simple cells of complicated fate. Biological Reviews. 2024;:brv.13162. 10.1111/brv.13162.

67. Ereskovsky AV, Tokina DB, Saidov DM, Baghdiguian S, Le Goff E, Lavrov AI. Transdifferentiation and mesenchymal-to-epithelial transition during regeneration in Demospongiae (Porifera). Journal of Experimental Zoology Part B: Molecular and Developmental Evolution. 2019;334:37–58. 10.1002/jez.b.22919.

68. Fritzenwanker JH, Saina M, Technau U. Analysis of *forkhead* and *snail* expression reveals epithelial–mesenchymal transitions during embryonic and larval development of *Nematostella vectensis*. Developmental Biology. 2004;275:389–402. 10.1016/j.ydbio.2004.08.014.

69. Thiery JP, Acloque H, Huang RYJ, Nieto MA. Epithelial-Mesenchymal Transitions in Development and Disease. Cell. 2009;139:871–90. 10.1016/j.cell.2009.11.007.

70. Lamouille S, Xu J, Derynck R. Molecular mechanisms of epithelial–mesenchymal transition. Nat Rev Mol Cell Biol. 2014;15:178–96. 10.1038/nrm3758.

71. Sasakura Y, Hozumi A. Formation of adult organs through metamorphosis in ascidians. Wiley Interdisciplinary Reviews: Developmental Biology. 2018;7:e304.

72. Adamska M. Differentiation and Transdifferentiation of Sponge Cells. In: Kloc M, Kubiak JZ, editors. Marine Organisms as Model Systems in Biology and Medicine. Cham: Springer International Publishing; 2018. p. 229–53. 10.1007/978-3-319-92486-1_12.

73. Bump P, Khariton M, Stubbert C, Moyen NE, Yan J, Wang B, et al. Comparisons of cell proliferation and cell death from tornaria larva to juvenile worm in the hemichordate *Schizocardium californicum*. EvoDevo. 2022;13:13. 10.1186/s13227-022-00198-1.

74. Gifondorwa DJ, Leise EM. Programmed Cell Death in the Apical Ganglion During Larval Metamorphosis of the Marine Mollusc *Ilyanassa obsoleta*. The Biological Bulletin. 2006;210:109–20. 10.2307/4134600.

75. Roccheri MC, Tipa C, Bonaventura R, Matranga V. Physiological and induced apoptosis in sea urchin larvae undergoing metamorphosis. Int J Dev Biol. 2004;46:801–6. 10.1387/ijdb.12382946.

76. Sato Y, Kaneko H, Negishi S, Yazaki I. Larval arm resorption proceeds concomitantly with programmed cell death during metamorphosis of the sea urchin *Hemicentrotus pulcherrimus*. Cell Tissue Res. 2006;326:851–60. 10.1007/s00441-006-0212-6.

77. Seipp S, Schmich J, Leitz T. Apoptosis – a death-inducing mechanism tightly linked with morphogenesis in *Hydractina echinata* (Cnidaria, Hydrozoa). Development. 2001;128:4891–8. 10.1242/dev.128.23.4891.

78. Borchiellini C, Degnan SM, Le Goff E, Rocher C, Vernale A, Baghdiguian S, et al. Staining and Tracking Methods for Studying Sponge Cell Dynamics. In: Carroll DJ, Stricker SA, editors. Developmental Biology of the Sea Urchin and Other Marine Invertebrates. New York, NY: Springer US; 2021. p. 81–97. 10.1007/978-1-0716-0974-3_5.

79. Leys SP, Larroux C, Gauthier M, Adamska M, Fahey B, Richards GS, et al. Isolation of *Amphimedon* Developmental Material. Cold Spring Harb Protoc. 2008;2008:pdb.prot5095. 10.1101/pdb.prot5095.

80. Larroux C, Fahey B, Adamska M, Richards GS, Gauthier M, Green K, et al. Whole-Mount In Situ Hybridization in *Amphimedon*. Cold Spring Harb Protoc. 2008;2008:pdb.prot5096. 10.1101/pdb.prot5096.

